# SARS-CoV-2 infects cells following viral entry via clathrin-mediated endocytosis

**DOI:** 10.1101/2020.07.13.201509

**Authors:** Armin Bayati, Rahul Kumar, Vincent Francis, Peter S. McPherson

**Author notes:** These two authors contributed equally to this study (Armin Bayati, Rahul Kumar). Address correspondence to: Dr. Peter S. McPherson, Department of Neurology and Neurosurgery Montreal Neurological Institute, McGill University 3801 University Street, Montreal, QC H3A 2B4 Canada.

## Abstract

With more than 51 million cases and 1.3 million deaths, and with the resulting social upheaval, the COVID-19 pandemic presents one of the greatest challenges ever to human society. It is thus vital to fully understand the biology of SARS-CoV-2, the causative agent of COVID-19. SARS-CoV-2 uses its spike glycoprotein to interact with the cell surface as a first step in the infection process. Using purified spike glycoprotein and lentivirus pseudotyped with spike glycoprotein, we now demonstrate that following engagement with the plasma membrane, SARS-CoV-2 undergoes rapid clathrin-mediated endocytosis. This suggests that transfer of viral RNA to the cell cytosol occurs from the lumen of the endosomal system, and importantly clathrin-heavy chain knockdown, which blocks clathrin-mediated endocytosis, reduces viral infectivity. This discovery reveals important new information about the basic biology of SARS-CoV-2 infectivity.

---

The internal membrane system represents a major evolutionary advance associated with the emergence of the eukaryotic cell. Its dynamics are integral to basic cellular function and underlie much of the novel capability of the cell, including the almost infinite range of physiological responsiveness required for the complexity of eukaryotic organisms. Unfortunately, viruses subvert this complexity to gain access into cells and to propagate in a manner that allows them to limit immune surveillance. Antiviral strategies targeting the earliest steps of infection, such as cellular entry are appealing, as interference through a common, early step can provide robust efficacy. Thus, understanding the mechanisms of viral cellular entry is crucial.

As of November 11th, 2020 there were over 51 million confirmed cases of COVID-19 worldwide with ~1.3 million deaths (coronavirus.jhu.edu). However, studies of antibody seroprevalence have suggested that between 3 and 20% of total populations have been exposed to severe acute respiratory syndrome coronavirus-2 (SARS-CoV-2), the COVID-19 causing virus (Bendavid et al., 2020; Goodman and Rothfeld, 2020; Anand et al., 2020). Thus, infection rates are higher than viral tests indicate, but well below a level that would begin to allow herd immunity. Despite the prevalence and worldwide impact of COVID-19 on human health and the global economy, many aspects of fundamental virus biology remain unknown.

Since the beginning of the 21^st^ century, 3 coronaviruses have crossed the species barrier to cause deadly types of pneumonia in humans; Middle-East respiratory syndrome coronavirus (MERS-CoV) (Zaki et al., 2012), SARS-CoV (Drosten et al., 2003; Ksiazek et al., 2003), and SARS-CoV-2 (Huang et al., 2020; Zhu et al., 2020). All originated in bats but zoonotic transmission involved intermediate hosts; dromedary camels (MERS-CoV), civets (SARS-CoV), and unknown (SARS-CoV-2). The presence of numerous coronaviruses in bats suggests that zoonotic transmission to humans will continue (Tortorici and Veesler, 2019), and new pandemic potential viruses continue to emerge (Sun et al., 2020).

SARS-CoV-2 is a single-stranded RNA virus in which a lipid membrane surrounds the RNA and nucleocapsid proteins. Other viral structural proteins associate with the viral membrane including the spike protein, a transmembrane glycoprotein that forms homotrimers (Tortorici and Veesler, 2019). The spike glycoprotein is composed of S1 and S2 subdomains (Wrapp et al., 2020). The S1 subdomain encodes the receptor binding domain and is responsible for binding to host cells, whereas the S2 subdomain encodes the transmembrane portion of the spike protein and is responsible for fusion of the viral membrane with cellular membranes.

The receptor for the spike glycoprotein is angiotensin converting enzyme 2 (ACE2) (Hoffmann et al., 2020; Letko et al., 2020; Zhou et al., 2020; Walls et al. 2019). ACE2 is a transmembrane metallopeptidase localized primarily on the plasma membrane of many cell types with abundant expression in lung alveolar epithelial cells (Hamming et al., 2004). The receptor binding domain of the spike protein binds ACE2 with low nM affinity, allowing the virus to stably associate with the plasma membrane (Walls et al., 2019). Cleavage of the spike glycoprotein between the S1 and S2 domains, mediated by the type II transmembrane serine protease TMPRSS2 (Hoffmann et al., 2020), and perhaps by furin, activates the S2 subdomain. The S2 subdomain then mediates the fusion of the viral membrane and the cellular membrane in a process that is a molecular mimic of SNARE-mediated fusion of cellular membranes (Walls et al., 2019; Hamming et al., 2004; Weber et al., 1998). This creates a pore allowing the RNA and RNA-associated nucleocapsid proteins within the lumen of the virus to gain access to the cellular cytosol, triggering infection.

For SARS-CoV-2, it remains unknown where the fusion of the viral and cellular membranes occurs. One possibility is that fusion occurs at the cell plasma membrane (Tang et al., 2020). In such a scenario, the virus does not directly enter the cell, but the viral RNA that drives infection enters the cytosol through a fusion pore in the plasma membrane. Alternatively, SARS-CoV-2 may undergo endocytosis with the entire virion particle rapidly entering the cell. In this scenario, the virion membrane would fuse with the luminal face of the endosomal membrane, allowing for RNA transfer to the cytosol. In fact it appears that most human coronaviruses are engulfed/endocytosed before they can infect the cell with their genetic material. This includes human (H)CoV-229E (Yeager et al., 1992; Nomura et al., 2004) HCoV-OC43 (Owczarek et al., 2018), HCoV-NL63 (Hofmann et al., 2005; Milewska et al., 2018), HCoV-HKU1 Hulswit et al., 2019), MERS-CoV (Burkard et al., 2014; Lu et al., 2013) and SARS-CoV (Inoue et al., 2007, Wang et al., 2020). And yet the endocytic mechanisms used by these coronaviruses remain unclear as they have been shown to use: 1) clathrin-dependent; 2) caveolae-dependent; 3) caveolae- and clathrin-independent pathways. In fact, the virus most similar to SARS-CoV-2 is SARS-CoV and there are conflicting results regarding its endocytic pathway; one study indicating that the virus uses clathrin-mediated endocytosis (Inoue et al., 2007) and a second stating the opposite, that viral entry prior to infectivity uses a clathrin-independent process (Wang et al., 2008). Here we demonstrate rapid clathrin-mediated endocytosis of SARS-CoV-2 and provide evidence that this process is critical for infectivity.

## Results

### SARS-CoV-2 spike glycoprotein is rapidly endocytosed in an ACE2-dependent manner

The spike glycoprotein is critical for binding ACE2 and allowing for SARS-CoV-2 infectivity (Tortorici and Veesler, 2019; Wrapp et al., 2020; Hoffmann et al., 2020; Letko et al., 2020; Zhou et al., 2020; Walls et al., 2020; Hamming et al., 2020; Weber et al., 1998). Wild-type HEK-293T cells or HEK-293T cells stably expressing ACE2 (Crawford et al., 2020) were incubated for 30 min on ice with purified spike protein containing a His6 tag. Following a PBS wash, spike protein binds to the plasma membrane of cells expressing ACE2 but not to the control HEK-293T cells (Fig. 1A), which have very low endogenous levels of the ACE2 receptor (Fig. 6A). In contrast, transferrin (Trf), which binds to the Trf receptor, is detected on the surface of the cells independent of their ACE2 status (Fig. 1A). Following a brief acid wash, both spike protein and Trf are stripped from the surface of cells (Fig. 1B). Thus, HEK-293T cells lacking or expressing ACE2 are a functional model system for examining potential SARS-CoV-2 endocytosis.

**Figure 1.**
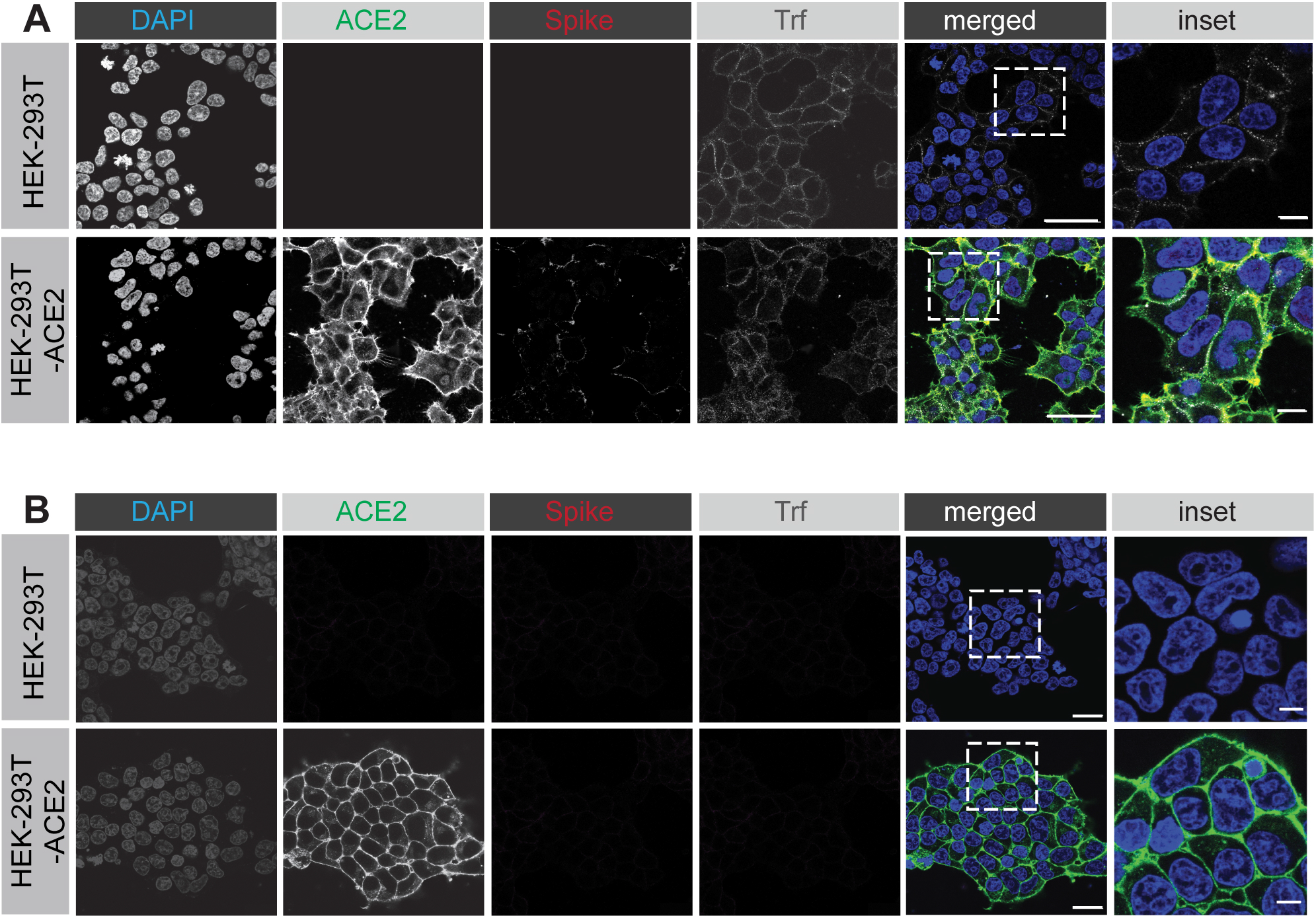
SARS-CoV-2 spike protein binds to the surface of HEK-293T cells expressing ACE2. **(A)** HEK-293T cells, wild-type (top row of images) or stably expressing ACE2 (bottom row of images) were incubated with purified, His6-tagged spike protein and with alexa-647 labelled transferrin(Trf) for 30 min at 4°C. Following PBS wash, the cells were fixed and stained with DAPI to reveal nuclei, with an antibody recognizing the expressed ACE2, and with an antibody recognizing the His6 epitope tag of the spike protein. Scale bars = 40 μm for the low mag images and 10 μm for the higher mag inset of the merged images. **(B)** Experiment performed as in **A** except that the HEK-293T cells were briefly acid washed prior to fixation. Scale bars = 40 μm for the low mag images and 10 μm for the higher mag inset of the composite.

We first tested for endocytosis using purified spike glycoprotein. While a reductionist approach, it is unlikely that receptor binding and initial membrane trafficking depends on whether or not the spike protein is on a viral particle. For example, epidermal growth factor and insulin both undergo similar cell surface binding and clathrin-mediated endocytosis whether they are purified proteins or attached to magnetic beads (Li et al., 2005). We thus added purified spike protein to HEK-293T cells, wild-type or expressing ACE2. Following 30 min on ice, the cells were transferred to 37°C for an additional 30 min. Following this treatment, cells were washed briefly with acid to strip off any surface bound spike protein. Spike protein was internalized into the cells (as revealed by its resistance to acid wash) in an ACE2-dependent manner (Fig. 2). In contrast, Trf was internalized independent of ACE2 (Fig. 2).

**Figure 2.**
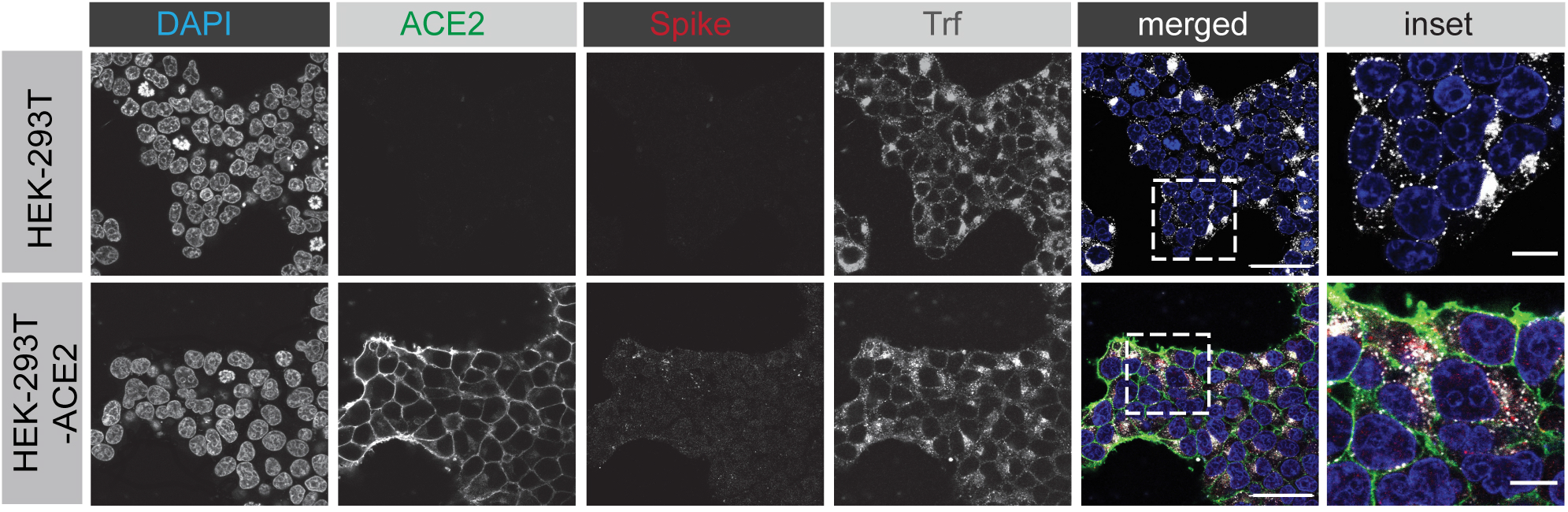
SARS-CoV-2 spike protein enters cells in an ACE2-dependent manner. HEK-293T cells, wild-type (top row of images) or stably expressing ACE2 (bottom row of images) were incubated with purified, His6-tagged spike protein and with alexa-647 labelled transferrin (Trf) for 30 min at 4°C. The cells were then transferred to 37°C for 30 min. The cells were returned to ice and following acid wash, were fixed and stained with DAPI to reveal nuclei, with an antibody recognizing the expressed ACE2, and with an antibody recognizing the His6 epitope tag of the spike protein. Scale bars = 40 μm for the low mag images and 10 μm for the higher mag inset of the composite.

We next examined the time course of internalization. Spike protein is internalized into cells rapidly and is detected in cells within 5 min (Fig. 3), a hallmark of endocytosis. The amount of spike protein in cells continues to increase for up to 45 min (Fig. 3). Thus, SARS-CoV-2 spike protein enters cells via endocytosis.

**Figure 3.**
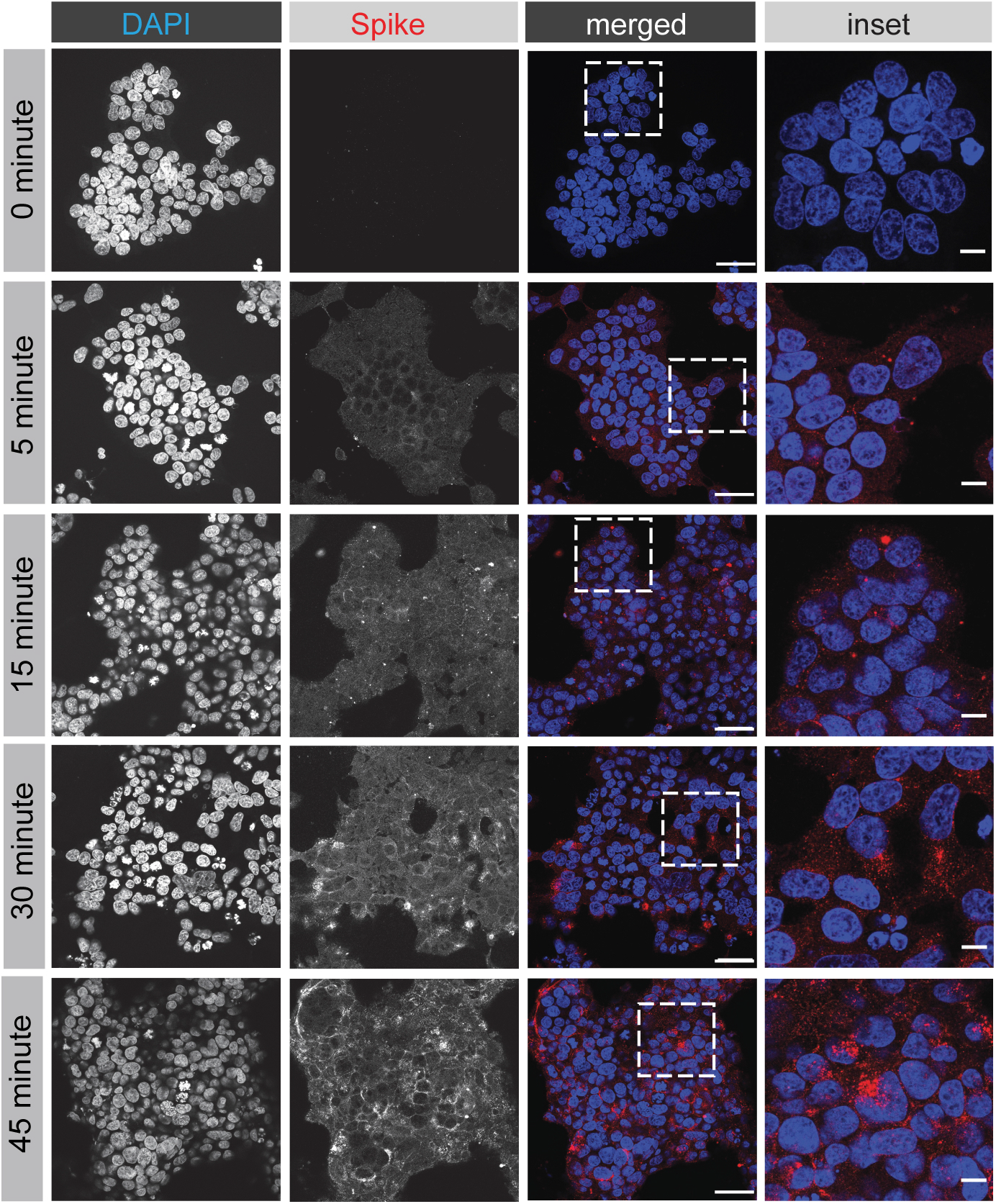
Time course of SARS-CoV-2 spike protein entry into cells. HEK-293T cells stably expressing ACE2 were incubated with purified, His6-tagged spike protein for 30 min at 4°C. The cells were then transferred to 37°C for the indicated time periods before being returned to ice. Following acid wash, cells were fixed and stained with DAPI to reveal nuclei and with antibody selectively recognizing the His6 epitope tag of the spike protein. Scale bars = 40 μm for the low mag images and 10 μm for the higher mag insets on the right.

### SARS-CoV-2 spike protein is internalized by clathrin-mediated endocytosis

There are conflicting reports about endocytosis of coronaviruses in general and SARS-CoV specifically (Inoue et al., 2007; Wang et al., 2008), and the endocytic mechanisms of SARS-CoV-2 have yet to be examined. Clathrin-mediated endocytosis is a major mechanism for cellular internalization. We thus used HEK-293T cells stably expressing ACE2 to examine the internalization of SARS-CoV-2 in the presence of two drugs that are known to block clathrin-mediated endocytosis, dynasore and Pitstop 2 (Marcia et al., 2006; von Kliest et al., 2011). Dynasore blocks the GTPase dynamin that drives the fission of clathrin-coated pits from the plasma membrane and Pitstop 2 prevents clathrin heavy chain (CHC) from interacting with adaptor proteins required for clathrin-coated pit formation (Marcia et al., 2006; von Kliest et al., 2011). Both drugs reduce the endocytosis of SARS-CoV-2 spike protein, strongly suggesting it is internalized by clathrin-mediated endocytosis (Fig. 4A/B). Both dynasore and Pitstop 2 have been criticized for having off-target effects (Park et al., 2013; Lemon and Traub, 2012). Therefore, we took a loss-of-function approach to inhibit clathrin-mediated endocytosis by knockdown of CHC with a previously defined siRNA pool (Galvez et al., 2007; Kim et al., 2011). Transfection of HEK-293T-ACE2 cells with the CHC siRNA led to an ~70% decrease in CHC levels when compared to cells transfected with a control siRNA (Fig. 5A). Importantly, CHC knockdown led to a significant reduction of spike protein endocytosis (Fig. 5B/C). Thus, the SARS-CoV-2 spike protein undergoes rapid internalization via clathrin-mediated endocytosis.

**Figure 4.**
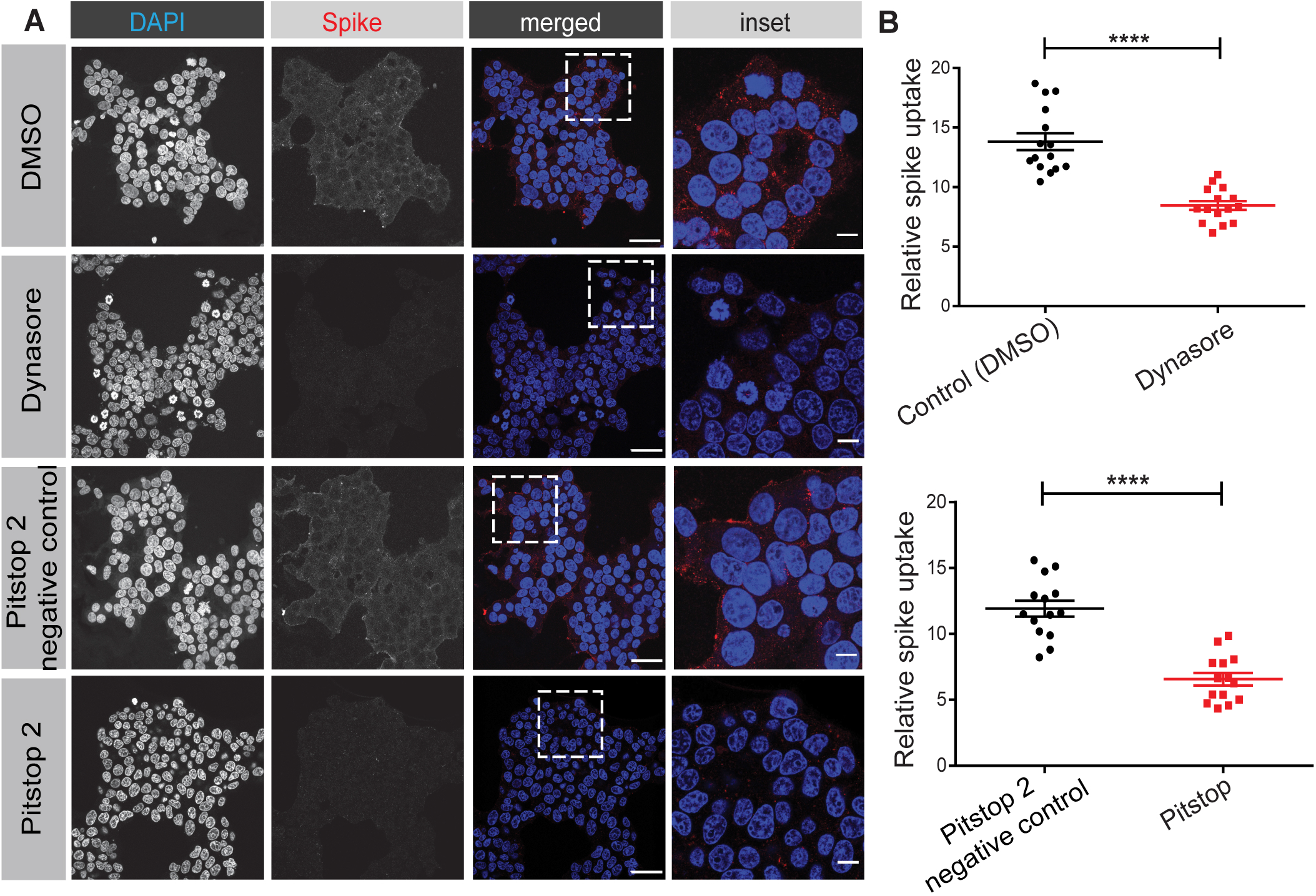
SARS-CoV-2 spike protein endocytosis is blocked by chemical inhibitors of clathrin-mediated endocytosis. **(A)** HEK-293 cells stably expressing ACE2 were incubated with purified, His6-tagged spike protein for 30 min at 4°C. The cells were then transferred to 37°C for 30 min before being returned to ice. Following acid wash, cells were fixed and stained with DAPI to reveal nuclei and with antibody selectively recognizing the His6 epitope tag of the spike protein. Inhibitors of clathrin-mediated endocytosis or their controls, as indicated, were added to the cells 30 min prior to the addition of spike protein. Scale bars = 40 μm for the low mag images and 10 μm for the higher mag insets on the right. **(B)** Quantification of experiments performed as in **A**. n=15 for Dynasore and n=14 for Pitstop 2 from three independent experiments, mean ± SEM; unpaired t-test; ****, p < 0.0001.

**Figure 5.**
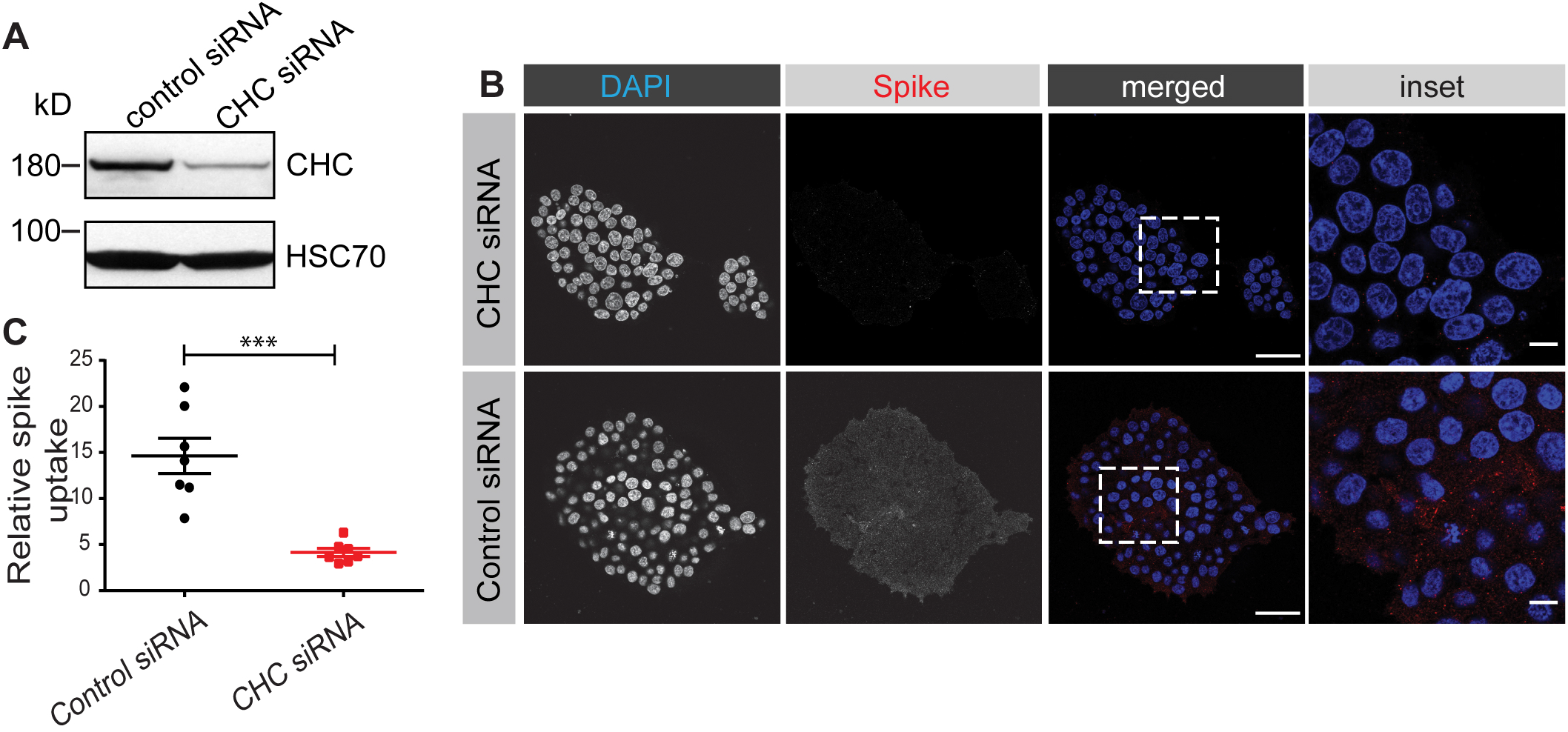
SARS-CoV-2 spike protein endocytosis is reduced by CHC knockdown. **(A)** HEK-293T cells stably expressing ACE2 were transfected with control siRNA or with an siRNA established to selectively knockdown the expression of CHC. Cell lysates were prepared and immunoblotted with the indicated antibodies. **(B)** HEK-293T cells stably expressing ACE2, transfected with a control siRNA or an siRNA driving CHC knockdown, were incubated with purified, His6-tagged spike protein for 30 min at 4°C. The cells were then transferred to 37°C for 30 min before being returned to ice. Following acid wash, cells were fixed and stained with DAPI to reveal nuclei and with antibody recognizing the His6 epitope tag of the spike protein. Scale bars = 40 μm for the low mag images and 10 μm for the higher mag insets on the right. **(C)** Quantification of experiments performed as in **B**. n=7 from three independent experiments, mean ± SEM; unpaired t-test; ***, p < 0.001.

### SARS-CoV-2 spike protein undergoes rapid endocytosis in cell lines expressing endogenous levels of ACE2

We next sought to examine endocytosis of SARS-CoV-2 spike protein in cells expressing endogenous levels of the cellular receptor ACE2. We first tested VERO cells, a monkey kidney epithelial cell that is robustly infected by SARS-CoV-2 (Hoffmann et al., 2020). VERO cells have high levels of ACE2 expression that exceeds even that of HEK-293T cells with overexpression of ACE2 (Fig 6A). SARS-CoV-2 spike protein added to VERO cells undergoes rapid endocytosis with the protein appearing in the cells as early as 5 min (Fig 6B), as determined using a spike protein antibody that recognizes the His6-tagged spike protein (Fig. S1). The amount of internalized protein increases slightly over the next 25 min with the protein appearing as distinct punctae in the cells (Fig. 6B). These punctate co-localize in part with Rab5, a marker of early endosomes, indicating that the spike glycoprotein is delivered to early endosomes (Fig. S2). Knockdown of CHC in VERO cells (Fig. 7A) reduces spike glycoprotein internalization (Fig. 7B/C), further supporting entry via clathrin-mediated endocytosis. We also tested for SARS-CoV-2 endocytosis in A549 cells that are widely used as a type II pulmonary epithelial cell model, and which are moderately infectible by the virus (Hoffman et al., 2020). A549 cells have ACE2 expression levels similar to the ACE2 expressing HEK-293T cell (Fig. 6A). These cells endocytose purified spike protein with the protein detectable in the cells at 5 min, and with increasing amounts of internalized protein up to 30 min (Fig. S3). As for the VERO cells, endocytosed spike protein appears as distinct punctae (Fig. S3). Thus, endocytosis of SARS-CoV-2 occurs in multiple cell types and is clathrin dependent.

**Figure 6.**
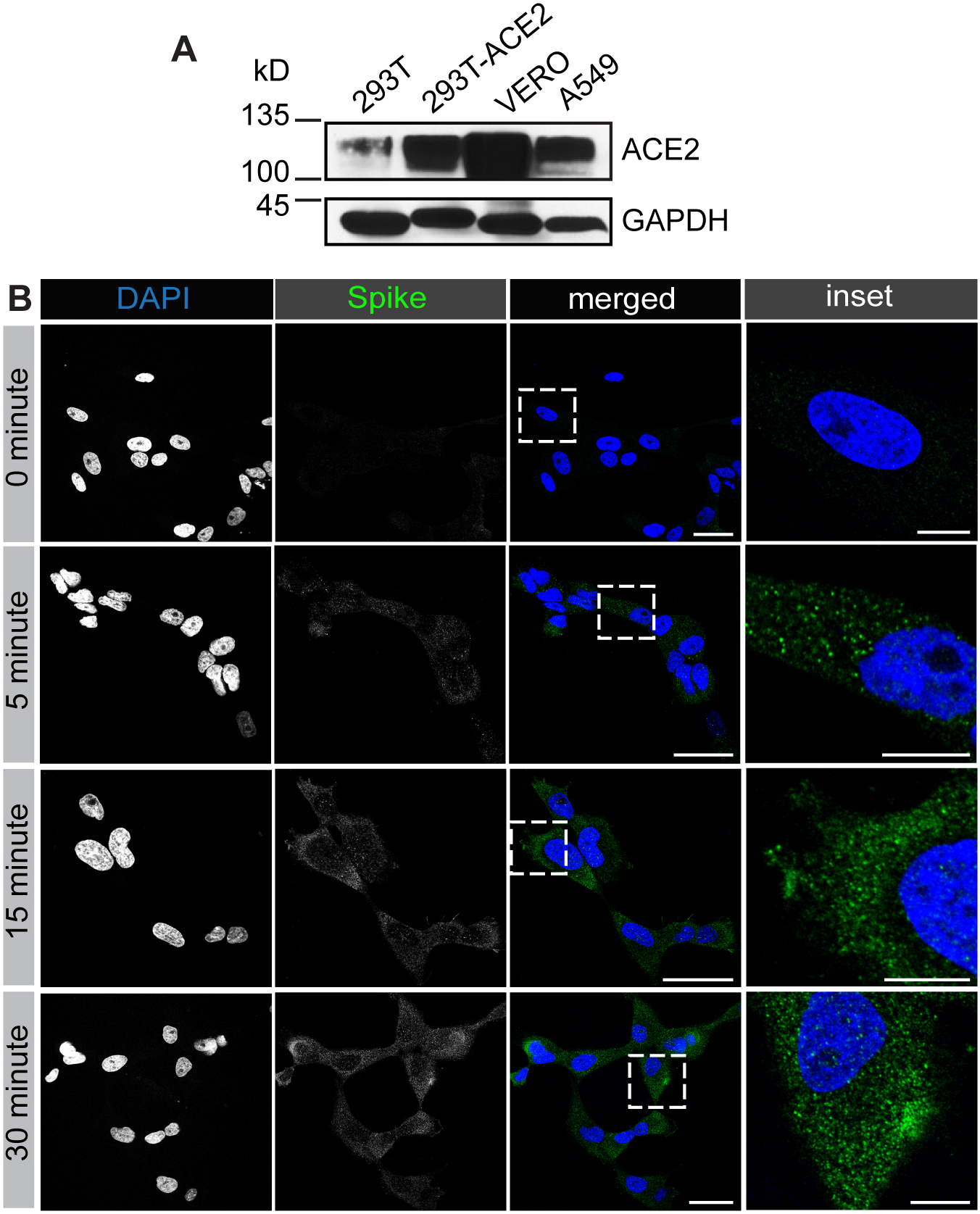
SARS-CoV-2 spike protein is rapidly endocytosed in VERO cells. **(A)** Lysates were made from HEK-293T cells, wild-type or expressing ACE2, VERO cells and A549 cells, and were immunoblotted with an antibody recognizing ACE2. **(B)** VERO cells were incubated with purified, His6-tagged spike protein for 30 min at 4°C. The cells were then transferred to 37°C for the indicated time periods before being returned to ice. Following acid wash, cells were fixed and stained with DAPI to reveal nuclei and with antibody selectively recognizing spike protein. Scale bars = 40 μm for the low mag images and 10 μm for the higher mag insets on the right.

**Figure 7.**
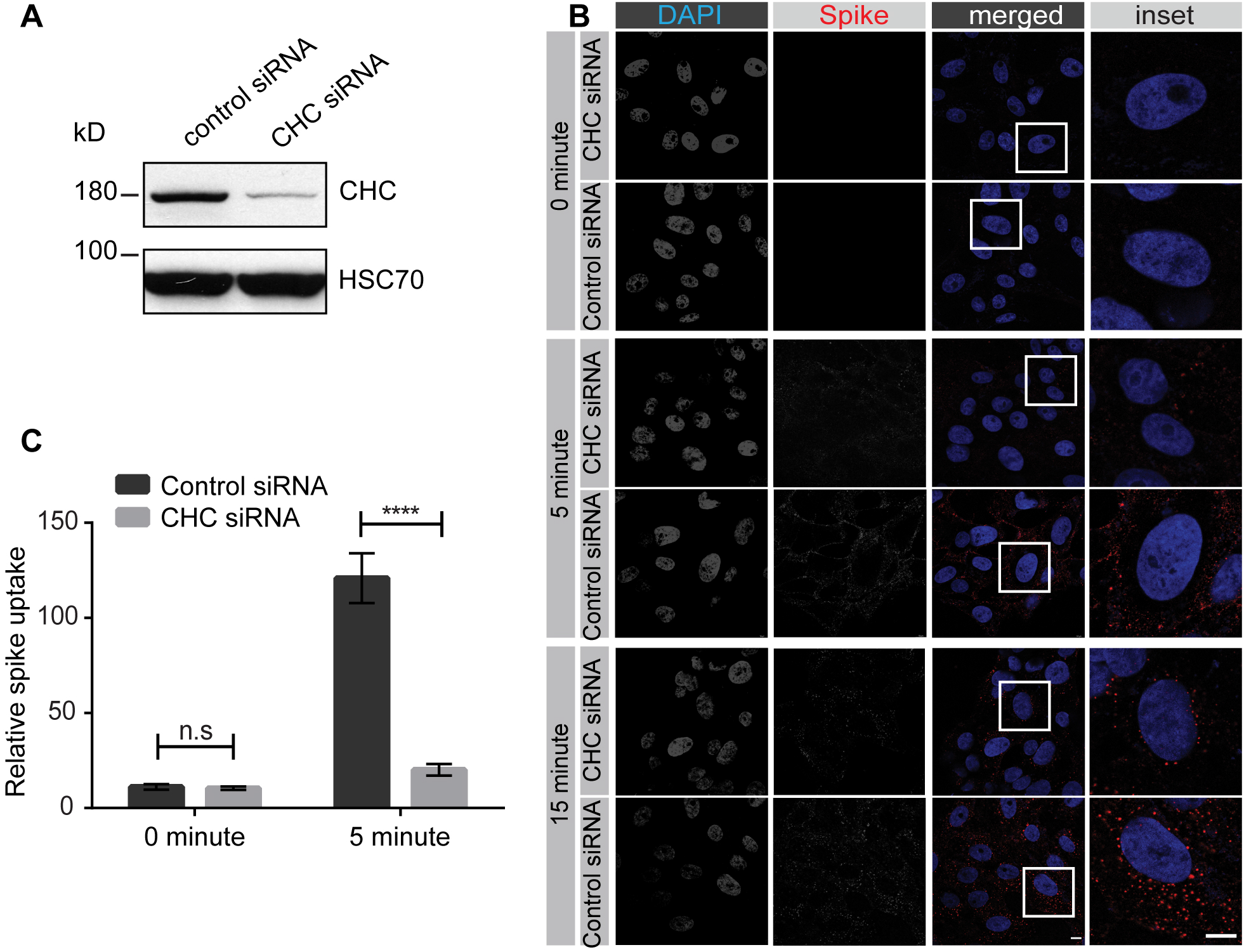
SARS-CoV-2 spike protein endocytosis is reduced by CHC knockdown in VERO cells. **(A)** Lysates were made from VERO cells transfected with control siRNA or with an siRNA established to selectively knockdown the expression of CHC. Cell lysates were prepared and immunoblotted with the indicated antibodies. **(B)** VERO cells, treated as in **A**, were incubated with purified spike protein for 30 min at 4°C. The cells were then transferred to 37°C for the indicated time periods before being returned to ice. Following acid wash, cells were fixed and stained with DAPI to reveal nuclei and with antibody selectively recognizing spike protein. Scale bars = 40 μm for the low mag images and 10 μm for the higher mag insets on the right. **(C)** Graph showing quantification of experiments performed as in **B**. n = 15 from 3 independent experiments, mean ± SEM; unpaired t-test; ****, p < 0.0001, n.s = not significant.

### Following endocytosis SARS-CoV-2 spike protein has a distinct trafficking itinerary from Trf

Trf bound to Trf-receptor is an established cargo of the recycling endosomal pathway. The Trf/Trf-receptor complex enters cells via clathrin-mediated endocytosis and rapidly transports to early endosomes. From there the complex recycles directly back to the plasma membrane, a fast recycling route, or is transported to Rab11-positive recycling endosomes before returning to the cell surface, the slow recycling pathway (Stoorvogel et al., 1991). When spike protein and transferrin are co-incubated with HEK-293T cells stably expressing ACE2, both Trf and spike protein are rapidly internalized and accumulate over 30 min (Fig. 8). However, there is little or no co-localization of the two cargo proteins once internalized (Fig. 8), indicating that they have different trafficking itineraries.

**Figure 8.**
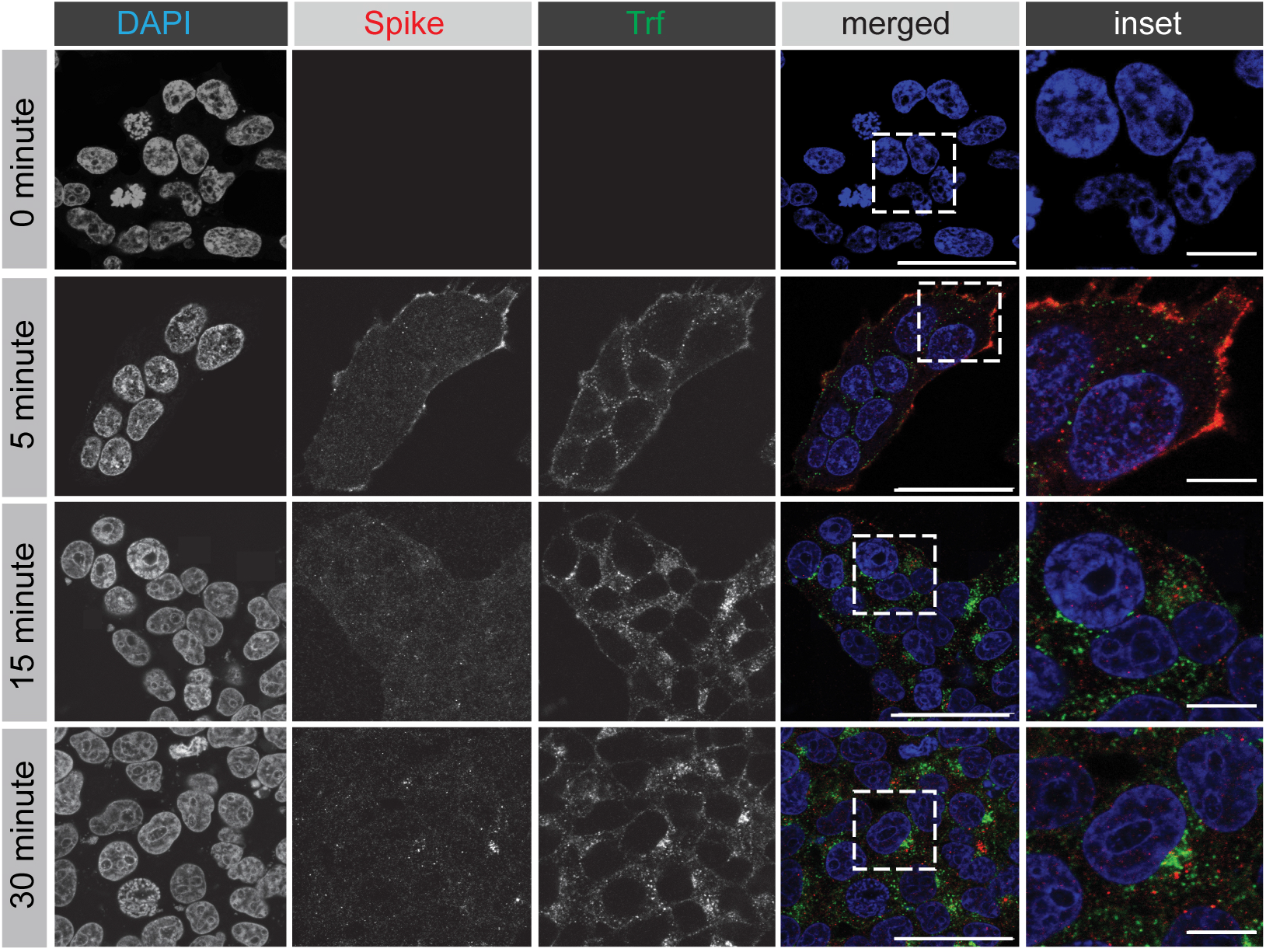
SARS-CoV-2 spike protein follows a trafficking itinerary distinct from transferrin. HEK-293T cells stably expressing ACE2 were incubated with purified, His6-tagged spike protein and Trf-Alexa647 for 30 min at 4°C. The cells were then transferred to 37°C for the indicated time periods before being returned to ice. Following PBS wash, cells were fixed and stained with DAPI to reveal nuclei and with antibody selectively recognizing the His6 epitope tag of the spike protein. Scale bars = 40 μm for the low mag images and 10 μm for the higher mag insets on the right.

### Lentivirus pseudotyped with SARS-CoV-2 spike glycoprotein requires clathrin for infectability

We next sought to examine the role of clathrin-mediated endocytosis in viral infectivity. Viruses pseudotyped with the SARS-CoV-2 spike glycoprotein are a commonly used tool to examine the ability of SARS-CoV-2 to infect cells. Such pseudotyped viruses involve transfection of HEK-293T cells with a viral packaging construct, a viral transfer vector encoding luciferase or GFP, and a plasmid encoding the spike protein, and are based on murine leukemia virus (Walls et al., 2020), VSV (Kang et al., 2020), or lentivirus (Crawford et al., 2020) among others. These pseudoviruses avoid the need for biocontainment 3 facilities. We have used a lentiviral system in which VSV-G was replaced with the SARS-CoV-2 spike glycoprotein (Crawford et al., 2020). We first incubated the virus with HEK-293T cells, wild-type or overexpressing ACE2, and infection was monitored by GFP expression. At 12 hours of incubation, HEK-293T-ACE2 cells were infected, whereas little to no infection was seen with wild-type HEK-293T cells (Fig. 9A). We next repeated the experiment using HEK-293T-ACE2 cells transfected with a control siRNA or an siRNA driving knockdown of CHC (Fig. 5A). Importantly, CHC knockdown led to an ~65% decrease in viral infectivity (Fig. 9B/C). Taken together, our data indicate that SARS-CoV-2 uses its spike glycoprotein to engage ACE2, driving clathrin-mediated endocytosis of the virus/receptor complex, and that this process is required for viral infectivity.

**Figure 9.**
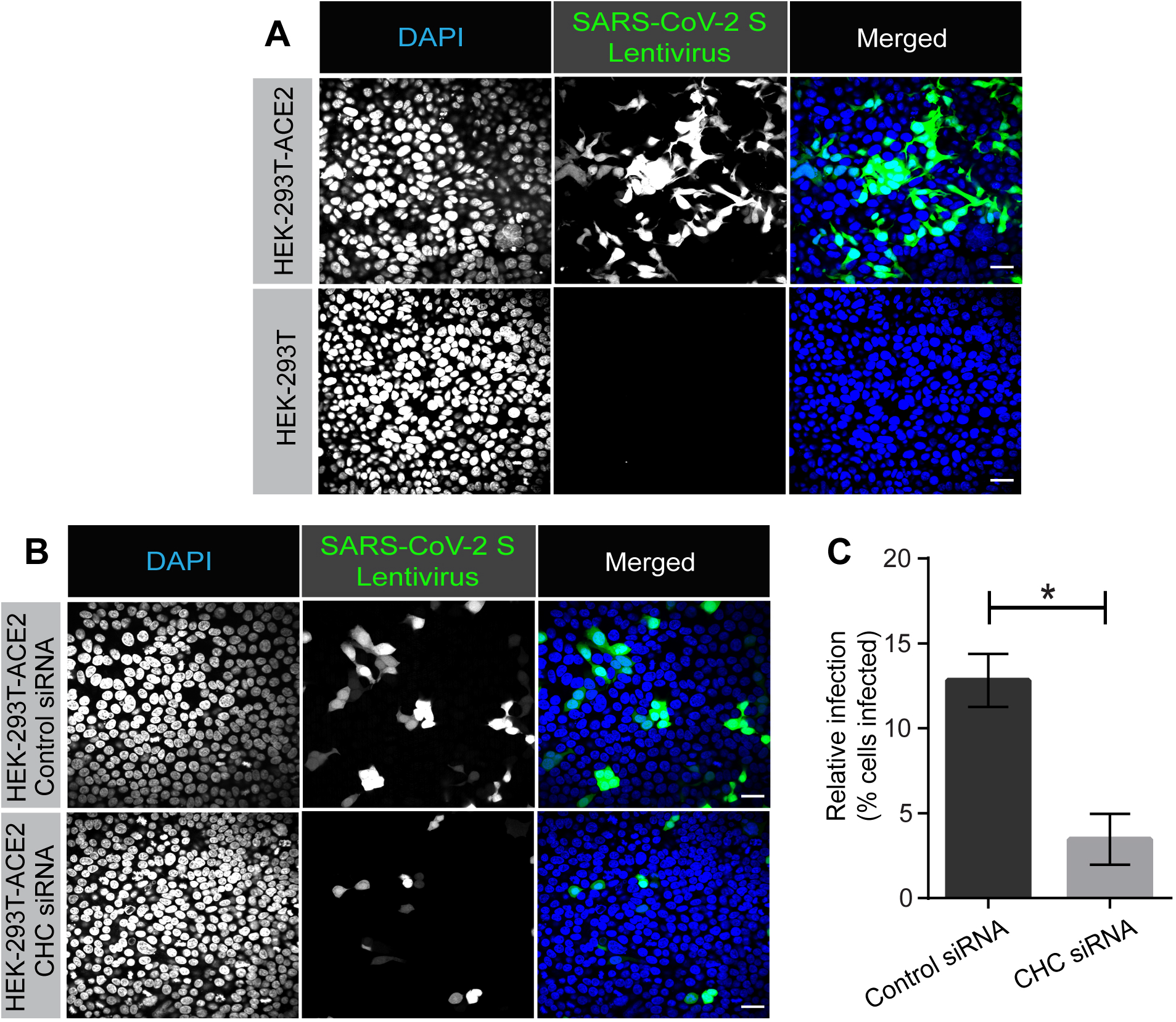
Lentivirus pseudotyped with SARS-CoV-2 spike glycoprotein requires CHC for infectivity. **(A)** HEK-293T cells stably expressing ACE2 or HEK-293T cells were incubated with lentivirus pseudotyped with the SARS-CoV-2 spike glycoprotein for 12 h. Cells were then fixed and stained with DAPI to reveal nuclei. Cells were also imaged for GFP, driven from the pseudovirus. **(B)** HEK-293T cells stably expressing ACE2 were transfected with control siRNA or with an siRNA established to selectively knockdown the expression of CHC. The cells were incubated with lentivirus pseudotyped with the SARS-CoV-2 spike glycoprotein for 12 h. Cells were then fixed and stained with DAPI to reveal nuclei. Cells were also imaged for GFP, driven from the pseudovirus. **(C)** Graph showing quantification of SARS-CoV-2 pseudovirus infection from experiments as in **B**. n = 3 from 3 independent experiments, mean ± SEM; unpaired t-test, *, p < 0.05.

## Discussion

Strategic entry mechanism used by viruses to infect mammalian host cells are unique and highly variable across different viral families (Pelkmans and Helenius, 2003). Among the various routes employed, the majority of viruses use endosomal pathways to efficiently deliver virion components into the cytoplasm for productive infection (Yamauchi and Helenius, 2013). While the most commonly used endosomal pathway is clathrin-mediated endocytosis (utilized for example by Semliki forest virus and Adenovirus 2 and 5 and), other forms of endocytosis are employed (Pelkmans and Helenius, 2003; Yamauchi and Helenius, 2013; Marsh and Helenius, 2006). These include caveolae-mediated endocytosis (e.g. Simian virus 40), lipid-raft-mediated endocytic pathways (e.g. Avian sarcoma leukosis virus), and macropinocytosis (e.g. poxviruses) (Pelkmans and Helenius, 2003; Yamauchi and Helenius, 2013; Marsh and Helenius, 2006).

Endosomal entry mechanisms provide many advantages to the viruses by allowing them to efficiently spread while evading host immune surveillance. Most importantly, gaining access to the endocytic system helps viruses to prevent exposure of the viral capsid proteins to the host immune system at the level of plasma membrane. Additionally, strategic use of endosomal pathway also ensures entry into cells with active membrane transport system as opposed to cells with no such means (e.g. erythrocytes), which would prevent active propagation of the virus (Pelkmans and Helenius, 2003). Further, the acidic milieu of endosomes helps the incoming viruses to elicit penetration into the host cytosol (Pelkmans and Helenius, 2003; Marsh and Helenius, 2006).

The most closely related virus to SARS-CoV-2 is SARS-CoV. For SARS-CoV, endocytic entry has been suggested as the first step in infectivity with two mechanisms proposed, one clathrin-mediated endocytosis (Inoue et al., 2007) with a second study indicating a clathrin-independent process (Wang et al., 2008). Here we used purified spike protein and lentivirus pseudotyped with SARS-CoV-2 spike glycoprotein to measure endocytosis and infectivity in HEK-293T cells expressing ACE2 and cell lines with endogenous ACE2 expression. We demonstrate that drugs, which are known inhibitors of clathrin-mediated endocytosis, block spike protein endocytosis and we further demonstrate that knockdown of CHC blocks both spike protein endocytosis and pseudovirus infectibility. We thus propose a viral infectivity model involving 3 key steps: 1) the virus uses its spike glycoprotein to bind to the plasma membrane of cells expressing ACE2. Notably, a recent paper questions that ACE2 is the cellular receptor for SARS-CoV-2 based on the observation that immunohistochemical analysis fails to detect ACE2 in cells and tissues that are infectible with virus (Hikmet et al., 2020). Our data does not support this observation; 2) the ACE2/virus complex undergoes rapid endocytosis with delivery to the lumen of the endosome; 3) fusion of the viral membrane with the lumen of the endosomal membrane allows viral RNA to enter the cytosol for infection. Following entry, the viral particles appear in a pathway that is unique from Trf. Since Trf defines a recycling pathway, we propose the viral capsid protein gets targeted to lysosomes for degradation.

The discovery that clathrin-mediated endocytosis is an early step in viral infectivity will allow for a new emphasis on drug targets. Chloroquine (CQ) and hydroxychloroquine (HCQ) are widely used malaria drugs that have yielded mixed results for the treatment of COVID-19 (Gao et al., 2020). CQ is known for its efficacy in blocking the clathrin-mediated endocytosis of nanoparticles (Hu et al., 2020). CQ reduces the expression of the phosphatidylinositol binding CHC assembly protein (PICALM) (Hu et al, 2020) and depletion of PICALM is known to block clathrin-mediated endocytosis (Miller et al., 2015). HCQ and CQ are in the class of aminoquinolines, which are hemozoin inhibitors, and were identified with other FDA-approved hemozoin inhibitors, including amodiaquine dihydrochloride dihydrate, amodiaquine hydrochloride and mefloquine, in a screen testing for reduction of SARS-CoV-2 infectivity (Weston et al., 2020). Chlorpromazine, which is widely used to treat psychiatric disorders, blocks cellular entry of SARS-CoV and is known to disrupt clathrin-mediated endocytosis (Inoue et al., 2007). This may explain why psychiatric patients treated with chlorpromazine have a lower incidence of COVID-19 (Plaze et al., 2020). Thus, while vaccine development is clearly the lead mechanism to slow the pandemic, drugs that disrupt virus infectivity are likely to have a corollary role. Regardless of clinical implications, given the prevalence and severity of SARS-CoV-2 infection, and the likelihood that other viruses with pandemic implications will emerge, it behooves the scientific community to learn about all aspects of SARS-CoV-2 infectivity, including basic functions such as its precise mechanism of cellular entry. This study, which provides a clear demonstration that clathrin-mediated endocytosis is employed by SARS-CoV-2 to enter cells, thus provides an important new piece of information on SARS-CoV-2 biology.

## Materials and Methods

### Cell lines

HEK-293T and A549 cell lines were obtained from ATCC (CRL-1573, CCL-185). Adherent Vero-SF-ACF cell line is from ATCC (CCL-81.5) (Kiesslich et. al., 2020). HEK-293T-ACE2 stable cell line was obtained from Dr. Jesse D. Bloom (Crawford et. al., 2020).

### Cell culture

All cells were cultured in DMEM high-glucose (GE Healthcare cat# SH30081.01) containing 10% bovine calf serum (GE Healthcare cat# SH30072.03), 2 mM L-glutamate (Wisent cat# 609065, 100 IU penicillin and 100 μg/ml streptomycin (Wisent cat# 450201). Cell lines were monthly checked for mycoplasma contamination using the mycoplasma detection kit (biotool cat# B39038).

### Immunoblot

Cells were collected in HEPES lysis buffer (20 mM HEPES, 150 mM sodium chloride, 1% Triton X-100, pH 7.4) supplemented with protease inhibitors. Cells in lysis buffer were gently rocked for 30 minutes (4°C). Lysates were spun at 238,700xg for 15 min at 4°C and equal protein aliquots of the supernatants were analyzed by SDS-PAGE and immunoblot. Lysates were run om large 5–16% gradient polyacrylamide gels and transferred to nitrocellulose membranes. Proteins on the blots were visualized by Ponceau staining. Blots were then blocked with 5% milk, and antibodies were incubated O/N at 4°C with 5% bovine serum albumin in TBS with 0.1% Tween 20 (TBST). The peroxidase conjugated secondary antibody was incubated in a 1:5000 dilution in TBST with 5% milk for 1 hr at room temperature followed by washes.

### Antibodies

SARS-Cov-2 spike protein antibody is from GeneTex (GTX632604). ACE2 antibody is from GeneTex (GTX01160). 6x-His Tag antibody is from ThermoFisher Scientific (MA1-21315-D550). GAPDH antibody is from OriGene (TA802519). HSC70 antibody is from Enzo (ADI-SPA-815-F). CHC antibody is from Cell Signaling (4796S). Conjugated Transferrin antibody is from ThermoFisher Scientific (T23366). Alexa Fluor 488, 568 and 647 conjugated secondary antibodies are from Invitrogen.

### Confocal microscopy

Cells were grown on poly-L-lysine coated coverslips. Cells were fixed in 4% PFA for 10 min and then washed 3 times with PBS. Cells were then permeabilized in 0.2% Triton X-100 in PBS and blocked with 2% BSA in PBS for 1 hour. Further, coverslips were incubated in a wet chamber with diluted antibody in blocking buffer overnight at 4°C. The following day, cells were washed 3 times and incubated with corresponding Alexa Fluorophore diluted in blocking buffer for 1 h at room temperature. Cells were again washed 3 times with blocking buffer and once with PBS. Nuclei were stained using DAPI (1 μg/ml diluted in blocking buffer) for 10 min. Finally, coverslips were mounted on a slide using mounting media (DAKO, Cat# S3023). Imaging was performed using a Leica TCS SP8 confocal microscope, Zeiss LSM-880 and Opera Phoenix High-Content Screening microscope.

### Endocytosis assay using purified spike protein

Cells were incubated at 37°C with serum free starvation media (DMEM) for 3 h to enhance ACE2 receptor expression. Prior to the addition of spike protein, cells were cooled to 4°C by being placed on ice. Spike protein was added to each well (3 μg per 200 μl of media) and incubated on ice for 30 min (to allow ligand attachment to the cell surface). Subsequently, cells were incubated at 37°C (to allow internalization) for indicated time points (transferrin was also added in some experiments in the same manner). Prior to fixation, cells were acid washed (washing off extracellular spike protein) or PBS washed for 1 min and rinsed with acid or PBS. Cells were then washed 3 times with PBS followed by fixation for 10 min at 4°C.

### siRNA mediated knockdown of CHC

Various cell types at 60% confluency were transfected with siRNA targeted against CHC (Dharmacon; SMARTpool: ON-TARGETplus; L-004001-01-0010) or control siRNA (Dharmacon; ON-TARGETplus CONTROL) using jetPRIME reagent. On day 3, cells were processed for immunoblot to investigate the effect of siRNA and parallelly, cells were infected with pseudovirus or used for spike endocytosis assays.

### Purified SARS-Cov-2 spike and pseudovirus production

Purified SARS-CoV-2 spike protein prefusion-stabilized ectodomain (C-term His tag, with furin cleavage site removed, trimerization stabilized) was produced by LakePharma (#46328). And, lentivirus pseudotyped with SARS-CoV-2 Spike Protein was supplied by Creative Diagnostics (catalog number COV-PS02).

### Treatments with chemical inhibitors

Cells were washed five times with serum free starvation media (DMEM) to remove any traces of serum. Further, cells were incubated with Dynasore (80 μM; Abcam, Ab120192) or Pitstop 2 (15 μM; Abcam, ab120687) in serum free media for 30 or 20 min, respectively. DMSO was used as a control for Dynasore and Pitstop 2 negative control (Abcam, ab120688). Following incubation, drug containing media was replaced with spike protein (3 μg per 200 μl of media) and further incubated for 5 min to allow internalization. Finally, cells were processed for immunofluorescence labeling.

### Pseudovirus SARS-Cov-2 infection

Cells were transfected with control siRNA or siRNA targeted against CHC (Dharmacon) using jetPRIME. Cells were split on day 2 and 10,000 cells were seeded in each well (96 well-plate; CellCarrier-96 Ultra Microplates, Perkinelmer). At 12-14 h after cell seeding, cells were incubated with concentrated pseudovirus SARS-Cov-2 for 12 h. Following this incubation, pseudovirus containing media was replaced with regular DMEM media and incubated further for 48 h. Cells were fixed with 4% PFA for 10 min and stained with DAPI.

### Quantification

Internalization of purified spike: Image J was used to measure the fluorescence intensity. The DAPI channel was used to count the number of cells per image. This was done by calculating the maxima using the “Find Maxima” function. The threshold was set at 150. This automated the way in which to count the cells. Then, the spike protein fluorescence corresponding to the cell count was calculated using the “measure” function. Fluorescence was then divided by cell count to calculate the fluorescence per cell in each condition. Infection of cells using pseudovirus SARS-Cov-2: GFP expressing cells in CellCarrier-96 Ultra Microplate were imaged using Opera Phoenix HCS (40x objective). The total number of GFP expressing cells in each condition was normalized to the total number of cells in the respective wells.

### Statistics

Graphs were prepared using GraphPad Prism 6 software. For all data, comparisons were made using student’s T-test. All data are shown as the mean +/− SEM with P < 0.05 considered statistically significant.

## Acknowledgments

We acknowledge the Neuro Microscopy Imaging Centre and Advanced BioImaging Facility at McGill University. We thank Dr. Jesse D. Bloom (University of Washington, Seattle) for HEK-293T-ACE2 stable cell line and Dr. Amine A. Kamen (Department of Bioengineering, McGill University) for Vero ATCC CCL 81.5 cell line. This work was supported by a grant from the Natural Sciences and Engineering Research Council to PSM. AB is supported by a studentship from the Parkinson’s Society of Canada. RK is supported by a studentship from ALS Canada. VF was supported by a fellowship from the Fonds de recherche du Quebec - Sante. PSM is a James McGill Professor and a Fellow of the Royal Society of Canada.

## Figure legends

**Supplemental Figure 1.**
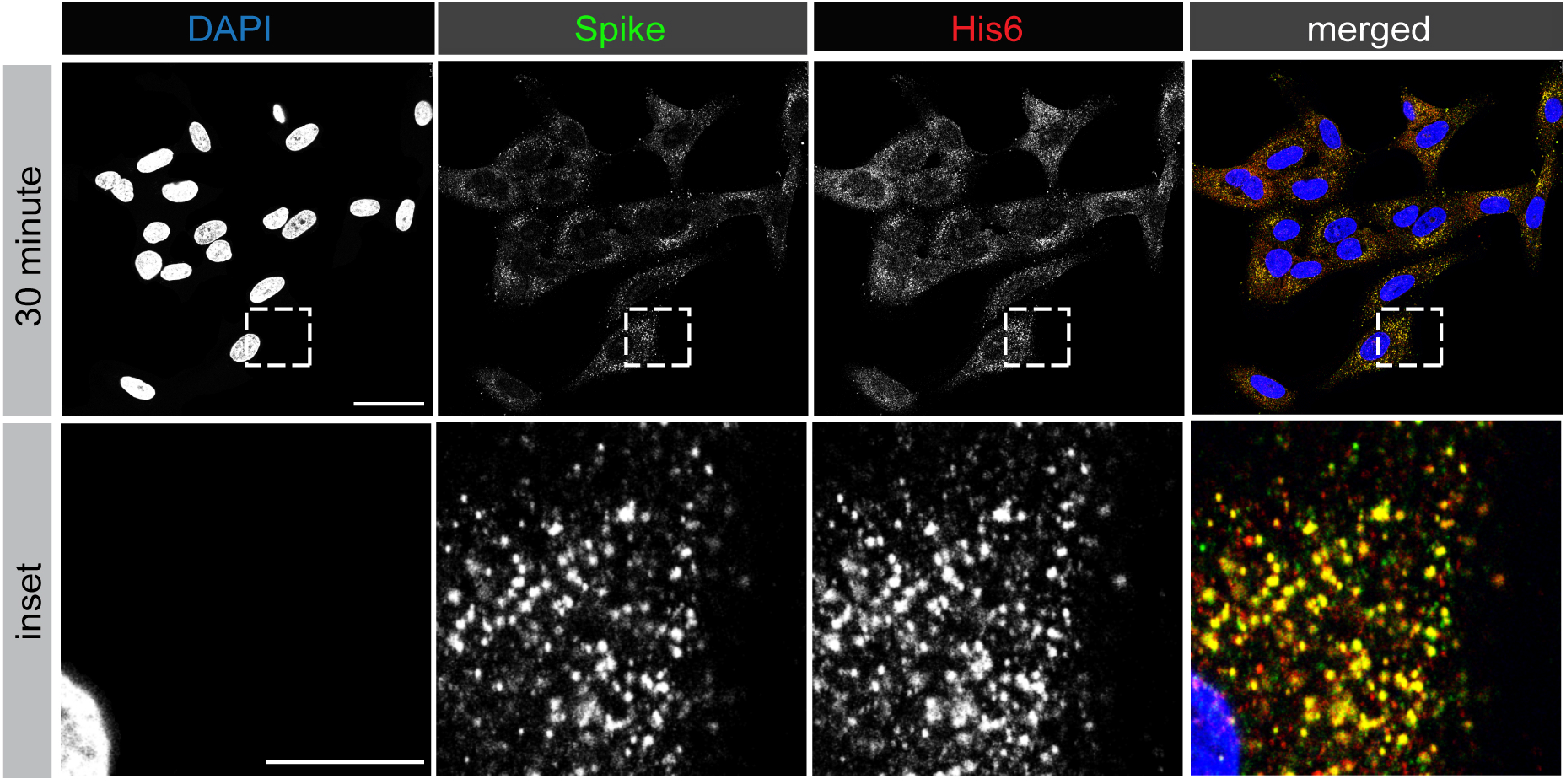
The antibody against SARS-CoV-2 spike protein specifically recognizes internalized protein. HEK-293T cells stably expressing ACE2 were incubated with purified, His6-tagged spike protein for 30 min at 4°C. The cells were then transferred to 37°C for the indicated time periods before being returned to ice. Following acid wash, cells were fixed and stained with DAPI to reveal nuclei and with antibody recognizing the His6 epitope tag of the spike protein and with antibody directed against the spike protein. Scale bars = 40 μm for the low mag images and 10 m for the higher mag insets on the right.

**Supplemental Figure 2.**
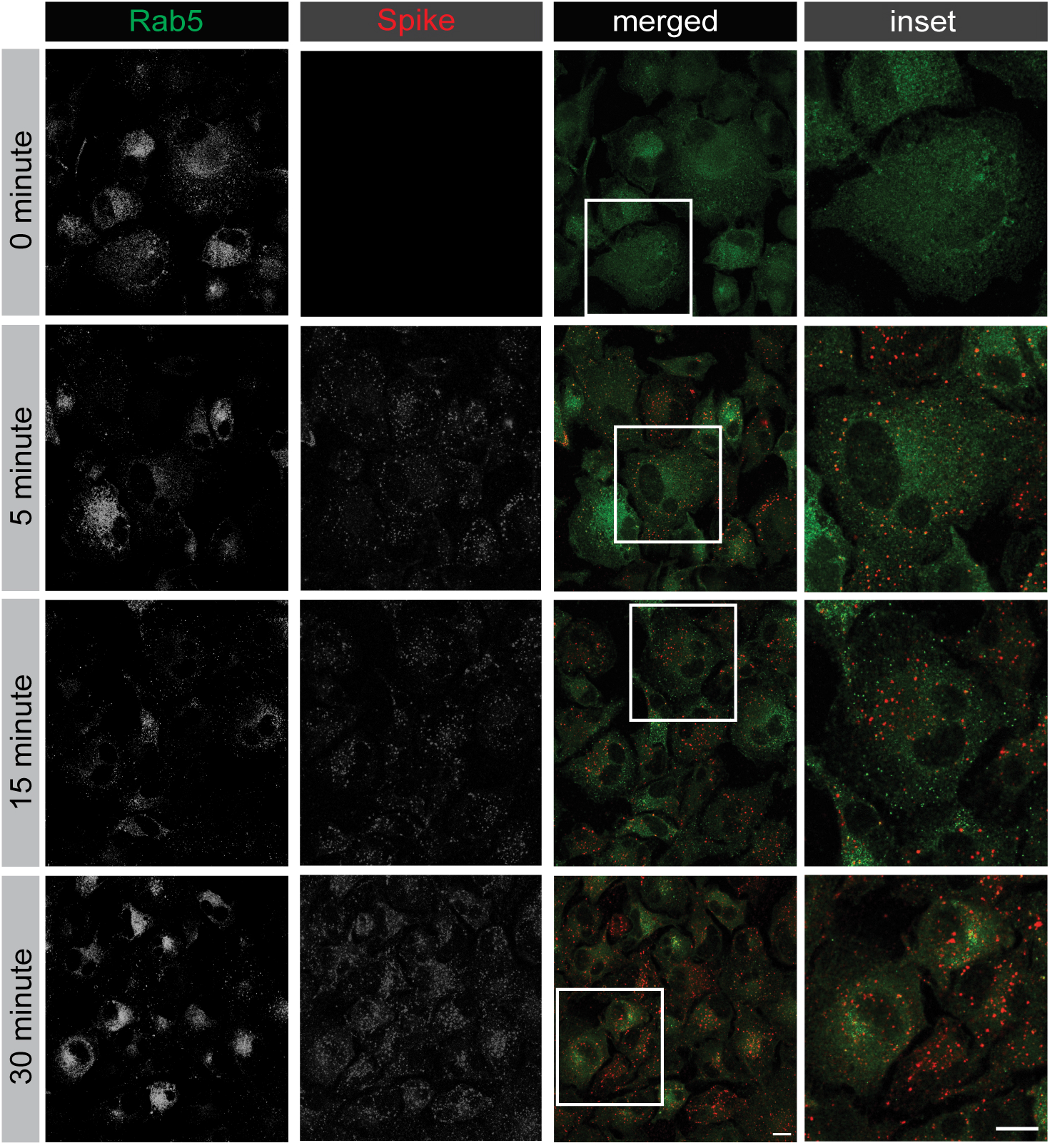
SARS-CoV-2 spike protein partially co-localizes with Rab5 following endocytosis. VERO cells were incubated with purified spike protein for 30 min at 4°C. The cells were then transferred to 37°C for the indicated time periods before being returned to ice. Following acid wash, cells were fixed and stained with antibody recognizing Rab5 and the spike protein. Scale bars = 40 μm for the low mag images and 10 μm for the higher mag insets on the right.

**Supplemental Figure 3.**
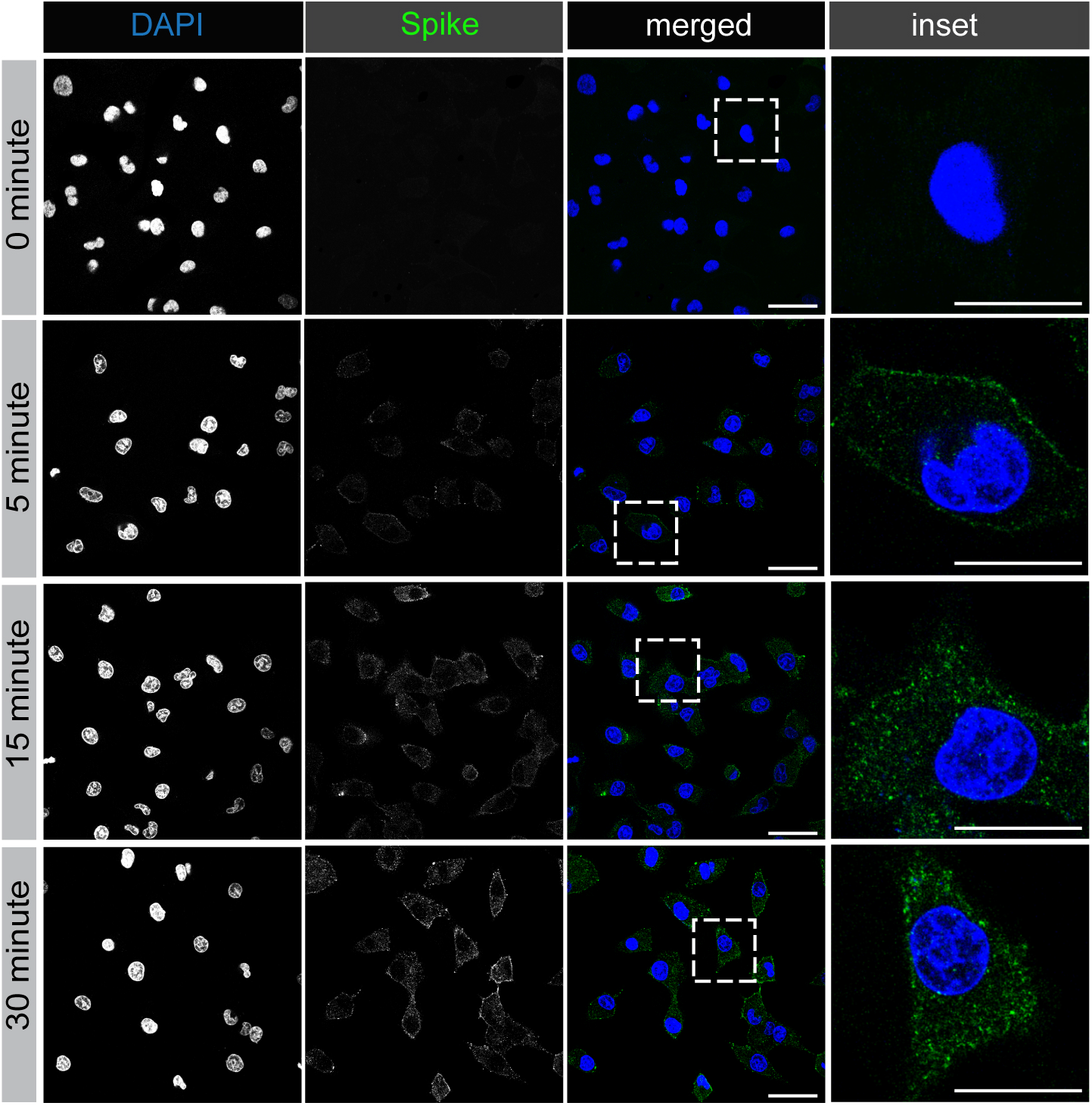
SARS-CoV-2 spike protein is rapidly endocytosed in A549 cells. A549 cells were incubated with purified, His6-tagged spike protein for 30 min at 4°C. The cells were then transferred to 37°C for the indicated time periods before being returned to ice. Following acid wash, cells were fixed and stained with DAPI to reveal nuclei and with antibody selectively recognizing spike protein. Scale bars = 40 μm for the low mag images and 10 μm for the higher mag insets on the right.

